# Prediction of meiosis-essential genes based on dynamic proteomes responsive to spermatogenesis

**DOI:** 10.1101/2020.02.05.936435

**Authors:** Kailun Fang, Qidan Li, Yu Wei, Jiaqi Shen, Wenhui Guo, Changyang Zhou, Ruoxi Wu, Wenqin Ying, Lu Yu, Jin Zi, Yuxing Zhang, Hui Yang, Siqi Liu, Charlie Degui Chen

## Abstract

Mammalian meiosis is a specific cell division process during sexual reproduction, whereas a comprehensive proteome of the different meiotic stages has not been systematically investigated. Here, we isolated different types of spermatocytes from the testes of spermatogenesis-synchronized mice and quantified the corresponding proteomes with high-resolution mass spectrometry. A total of 8,002 proteins were identified in nine types of germ cells, and the protein signatures of spermatogenesis were characterized using the dynamic proteomes. A supervised machine learning package, FuncProFinder, was developed to predict meiosis-essential candidate genes based on changes in their protein abundance. Of the candidates without functional annotation, four of the ten genes with the highest prediction scores, *Zcwpw1, Tesmin, 1700102P08Rik*, and *Kctd19*, were validated as meiosis-essential genes using knockout mouse models. The proteomic analysis of spermatogenic cells provides a solid foundation for studying the mechanism of mammalian meiosis.

## INTRODUCTION

Meiosis is a cell division process specific to germ cells, in which DNA replicates once and divides twice to generate four gametes. It is accepted that mammalian meiosis is a complex process including several molecular events as homologous recombination, synapsis and so on, but the genes involved in this process are not well characterized^1, 2^. In contrast, the genes participating in yeast meiosis are well studied, thus homology comparisons to yeast is a common strategy to investigate the meiotic genes in mammals (e.g., *Spo11, Dmc1, Psmc3ip*, and *Rnf212*)^3-7^. However, because the regulatory mechanism of mammalian meiosis is more complicated than that of yeast, the genes that specifically participate in mammalian meiosis cannot be found using this strategy. Knock-out of genes with testis- or oocyte-specific expression patterns is another approach to identify meiosis-essential genes. For instance, 54 testis-specific genes were knocked out by Miyata’s group, however, none of the knockout mice exhibited a meiosis-essential phenotype^8^. An efficient approach to find mammalian meiosis-essential genes is thus clearly required.

Since the status of gene expression is a fundamental characteristic tightly associated with physiological functions, a dynamic atlas of gene expression throughout spermatogenesis would be extremely useful for exploring meiosis-essential genes. Up to now, transcriptional gene expression in thousands of germ cells covering various developmental stages of spermatogenesis has been quantified at the single cell level^9-16^, resulting in a very detailed transcriptome landscape throughout spermatogenesis. Although 90% of protein-coded transcripts during spermatogenesis have been identified, this widespread transcription was suggested as a way to scan and correct DNA mutations in the germline^16^, and there was limited evidence indicating these transcripts could be translated to functional proteins in transcriptome studies. Additionally, multiple studies identified a poor correlation between mRNA and protein abundance in testes^17, 18^, and therefore, a global proteomic profiling of gene expression in spermatogenesis is of great meaning to unravel functional molecules in meiosis. However, reports regarding systematic profiling of proteomes during meiosis are limited. Only one type of meiotic cell – pachytene spermatocytes – was quantified in previous proteomics studies^7, 18^, leaving protein expression unknown in most stages of meiosis. Hence a comprehensive proteomic profiling of different cell types during mammalian meiosis is required here.

Although omics data can provide systematical views of biological process, how to identify functional genes from the datasets are difficult. Differentially expressed gene (DEG) analysis has been used to identify mammalian meiotic genes from omics data. For instance, a single-cell transcriptomic study of spermatogenesis successfully selected *Fbxo47* from a group of 1,223 leptotene or zygotene-abundant DEGs and verified that *Fbxo47* was meiosis-essential by gene knockout mice^9^. But how to select *Fbxo47* from the large pool of DEGs was of challenge. Recently, supervised machine-learning approaches were applied to systematically predict functional genes. For instance, autism risk genes were predicted by a supervised Bayesian method based on the differences of gene co-expression networks between the well-known autism genes and non-autism genes, and several predicted candidates were proofed to be within frequent autism-associated copy-number variants, implying the prediction could be true^19^.

In this work, to understand the molecular basis of mouse meiosis and predict meiosis-essential proteins, seven consecutive types of meiotic cells plus pre-meiotic spermatogonia and post-meiotic round spermatids were isolated, and the proteins in each cell-type were identified and quantified by high-resolution mass spectrometry using a label-free mode. The meiosis-dependent signatures were characterized by protein abundance changes. Furthermore, a supervised ensemble machine learning package, FuncProFinder, was developed to predict the meiosis-essential candidates, and five candidates - *Pdha2, Zcwpw1, Tesmin, Kctd19*, and *1700102P08Rik* - were verified as meiosis-essential genes using knockout mice. These comprehensive proteomics data paves a path to efficiently discover meiosis-essential proteins and to figure out their functions in meiosis.

## RESULTS

### Isolation of mouse spermatogenic cells

To quantify changes in protein abundance and closely monitor the molecular events in response to mouse meiosis, we isolated the spermatogenic cells before, during, and after meiosis in C57BL/6 mouse testes, including pre-meiotic TypeA undifferentiated spermatogonia, consecutive types of meiotic cells, and post-meiotic round spermatids.

The isolation workflow of spermatogenic cells is illustrated in Fig. 1a. TypeA undifferentiated THY1+ c-KIT-spermatogonia (Aundiff) were isolated from the testes of postnatal day 7 (P7) mice using magnetic activated cell sorting (MACS) according to an established method^20^ (Supplementary Information, Fig. S1a, c). The immuno-fluorescence staining of *PLZF*, a well-known Aundiff marker, revealed that the percentage of *PLZF*+ cells increased from 10% to 70% after purification (Table 1; Supplementary Information, Fig. S1b, d), implying that Aundiff cells were greatly enriched. The haploid round spermatids (RS) were purified by DNA-content–based cell sorting from the testes of P28 mice (Supplementary Information, Fig. S1e, g). DAPI-staining images indicated that the purity of isolated RS almost reached 100% (Table 1; Supplementary Information, Fig. S2f-i).

**Table 1.** The purity of the separated spermatogenic cells of different stages around meiosis

| Cell type | Mouse Num. | Cell Num. (×10<sup>6</sup>) | Cell Purity |  | Cell type | Mouse Num. | Cell Num. (×10<sup>6</sup>) | Cell Purity |  |
| --- | --- | --- | --- | --- | --- | --- | --- | --- | --- |
| earlyL | 9 | 4.99 | earlyL | 95.36% | midP | 6 | 5.78 | midP | 97.99% |
|  |  |  | L | 4.64% |  |  |  | earlyP | 2.01% |
| L | 5 | 4.34 | L | 78.57% | lateP | 9 | 21.16 | lateP | 96.62% |
|  |  |  | earlyL | 21.43% |  |  |  | earlyD | 3.38% |
| Z | 7 | 7.04 | Z | 88.40% | earlyD | 3 | 6.63 | earlyD | 90.59% |
|  |  |  | earlyP | 11.60% |  |  |  | lateP | 9.41% |
| earlyP | 8 | 6.06 | earlyP | 93.64% | lateD | 7 | 7.27 | lateD | 96.38% |
|  |  |  | Z | 6.36% |  |  |  | Diak. | 3.62% |
| Aundiff | 96 | 2.97 | PLZF+ | 79.38% | RS | 10 | 20.85 | RS | 100.00% |
|  |  |  | PLZF- | 20.62% |  |  |  |  |  |

**Fig. 1.**
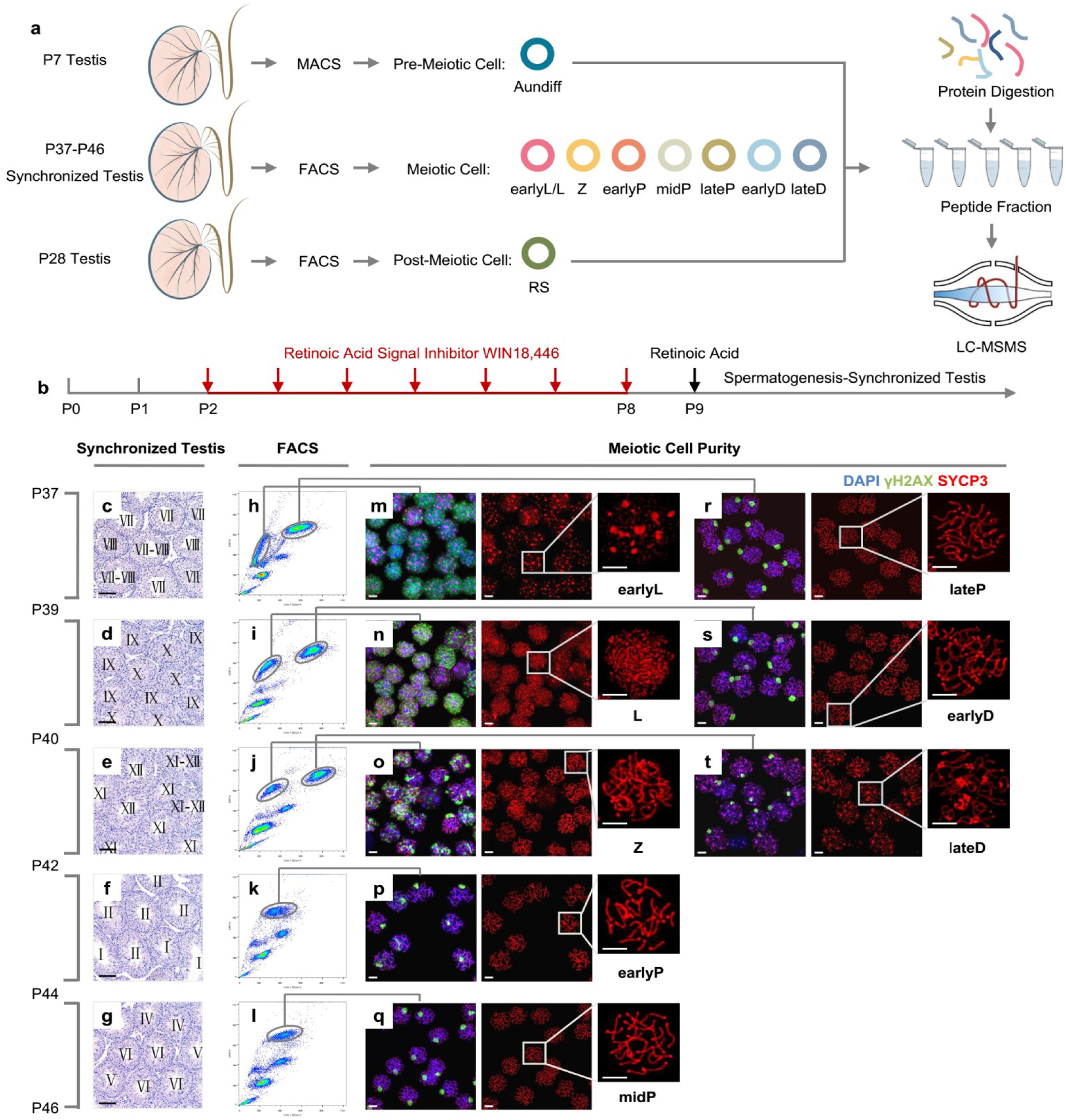
Isolation of spermatocytes during meiotic prophase I. **a** Workflow from germ cell collection to mass spectrometry analysis. **b** The strategy to obtain mice with synchronized spermatogenesis. **c-g** Cross sections of hematoxylin and eosin (H&E) stained testes from spermatogenesis-synchronized mice on Postnatal Day 37 (P37) to P46. Roman numerals in each tubule designate the stage represented by the cellular constitution with the criteria described previously^39^. Scale bar = 50 µm. **h-l** FACS plots of Hoechst 33,342 stained testes cells from the mice with synchronized spermatogenesis to the corresponding stages in c-g. **m-t** Chromatin spreading of FACS-sorted spermatocytes, that were co-immuno-stained with DAPI (blue), anti-γH2AX (green) and anti-SYCP3 (red). Scale bar = 5 µm.

In the seminiferous tubules of mouse testes, consecutive types of meiotic cells are mixed and are difficult to separate. To simplify the types of spermatocytes in testes, we applied a spermatogenesis synchronization method described before^9, 21, 22^: mouse spermatogonia differentiation was inhibited by WIN18,446 for seven days, and was re-activated synchronously by retinoic acid (RA) injection on P9 (Fig. 1b). Four weeks after RA treatment, testes of P37 to P46 mice exhibited only one or two types of meiotic spermatocytes at a given point (Fig. 1c-g), greatly facilitating DNA-content based cell sorting for purification (Fig. 1h-l). To assess the purity of the isolated meiotic cells, we performed immuno-fluorescence staining with antibodies against the synaptonemal complex (SC) marker *SYCP3* and the DNA damage marker γH2AX (Fig. 1m-t) and recognized spermatocyte cell-types with criteria described previously^23^. Based on the quantitative evaluation of fluorescence, most of isolated meiotic cells were of high purity at around 90% (Table 1). Considering that the protein amount of isolated early leptotene and leptotene were less than 120 μg, we mixed these two adjacent cell types together as an earlyL/L group for the following proteomic analysis. Thus, a total of seven types of meiotic cells, early leptotene and leptotene (earlyL/L), zygotene (Z), early pachytene (earlyP), middle pachytene (midP), late pachytene (lateP), early diplotene (earlyD), and late diplotene (lateD) were prepared for further proteomic study.

### Quantitative proteomic atlas of mouse meiosis during spermatogenesis

In the nine types of spermatogenic cells isolated above, a total of 8,002 proteins were identified (unique peptides≥2), with between 6,000–7,000 proteins in each cell type (Supplementary Information, Table S1). Within the identified proteins, 7,742 were only detected in seven sub-stages of meiosis, and 5,108 proteins were globally identified over all nine cell types. To obtain high quality quantification data, each sample was triplicated in LC-MS/MS. The Pearson correlation coefficients for all the triplicates in the same sample reached around 0.99 (Fig. 2a), indicating that proteomic quantification was highly replicated. Additionally, the comparison of protein expression correlation among the nine different cell-types revealed that the protein abundance changed dramatically between earlyP and midP, which strongly implied that spermatocytes undergo a cell state transition after passing of the midPachytene checkpoint.

**Fig. 2.**
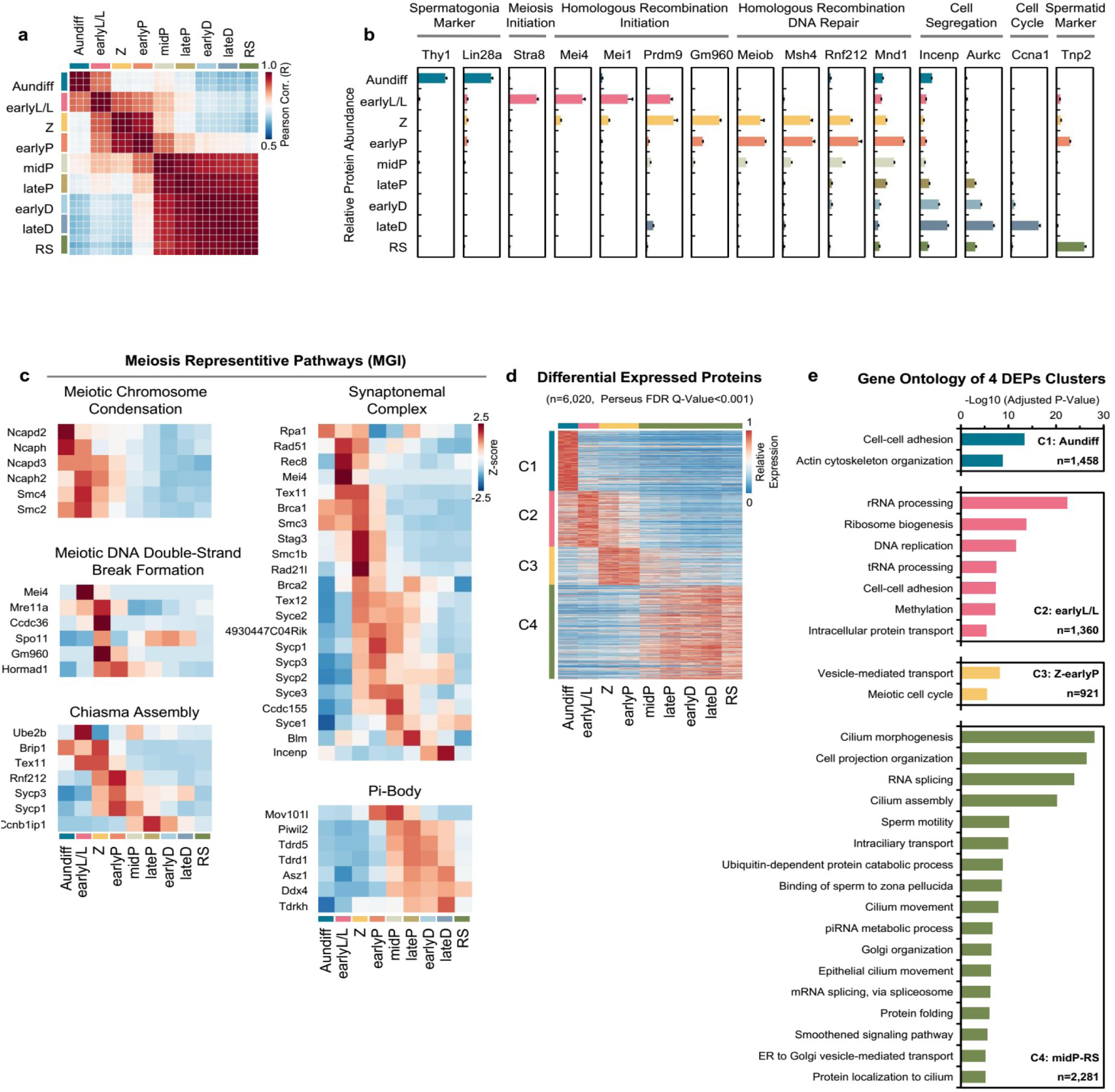
Proteomic informatics of mouse germ cells around meiotic prophase I. **a** Pearson correlation coefficients of the protein LFQ intensity of all quantified proteins among nine stages of germ cells. Color key represents the value of Pearson correlation coefficient. **b** Relative abundance of the typical biomarkers of germ cells in response to spermatogenesis. Error bars represent SEM in triplicates. **c** Heatmap of dynamic abundance of the proteins involved in five representative meiotic pathways (MGI database). **d** K-means clusters of relative protein abundance elicited from the DEPs. The color key represents the relative protein abundance. **e** Gene ontology (GO) analysis of the enriched biological processes for four DEP clusters.

Next, to evaluate the consistency of our protein quantification with previous knowledge, we tracked the protein abundance changes of 15 well-known cell-type-specific biomarkers throughout spermatogenesis (e.g., *LIN28A, STRA8, SPO11, TNP2* etc.). Protein abundances of these biomarkers appeared to be typically phase-dependent (Fig. 2b), which was basically in agreement with previous studies^24-27^. In addition, proteins in several meiosis-related processes, such as meiotic DNA double-strand break formation (MGI GO: 0042138), chiasma assembly (MGI GO: 0051026), synaptonemal complex (MGI GO: 0000800, 0000801, 0000802), are generally recognized as meiotic-phase dependent, and the proteomic evidence in this study further implied their functions (Fig. 2c). For instance, the SC mainly formed from Z to lateP and decreased after lateP^1^, and most SC components in proteomics data were consistent with previous results. However, protein abundance of *SYCE1*, a key SC component, remained stable from lateP to RS. A similar *Syce1* transcriptional expression pattern was observed from a single-cell RNA transcriptome dataset^9^, implying that *SYCE1* might perform additional functions except synapsis after meiosis Prophase I.

To further explore the phase-dependent dynamic processes, the abundance of all 8,002 identified proteins in nine cell types were analyzed using Perseus^28^ to identify differentially expressed proteins (DEPs). A DEP was defined as having significant changes in abundance between any two sub-stages when the Q-value was less than 0.001 with two-tailed test. A total of 6,020 proteins were determined to be DEPs, and these DEPs were divided into four groups by K-means analysis: C1 matched with Aundiff, C2 with earlyL/L, C3 with Z-earlyP, and C4 matched with midP-RS (Fig. 2d; Supplementary Information, Table S2). Gene ontology (GO) analysis towards DEPs in each cluster revealed the biological processes enriched in different phases (Fig. 2e). Cell-cell adhesion and actin cytoskeleton organization were enriched in Aundiff cells. Nucleic acid related processes such as rRNA processing and DNA replication were enriched in earlyL/L cells. Genes involved in meiotic cell cycle were enriched in Z-earlyP cells. piRNA metabolism, processes related to sperm functions, were enriched in midP-RS cells. The enriched KEGG pathways towards the four DEP clusters were also analyzed (Supplementary Information, Fig. S2a), and protein expressions of the four most significant-enriched pathways – DNA replication (C2 phase-dependent), spliceosome, proteasome, and oxidative phosphorylation (C4 phase-dependent) – were dynamically regulated during meiosis (Supplementary Information, Fig. S2b-e).

Taking all the information above, the high qualified dynamical proteomic data during spermatogenesis not only provided a bird’s eye view to categorize the meiosis progression at a global proteomic level, but also served as a resource to reveal previously uncharacterized molecular signatures of defined gene sets such protein complexes and even offered probability to identify additional functions of known meiotic protein.

### Prediction of meiosis-essential candidates using supervised machine learning analysis based on proteomic data

Although 8,002 proteins were identified in spermatogenesis and were further divided into four groups with relevant biological processes, the proteins essential to meiosis were still unclear. Recently, supervised machine-learning approaches were applied to systemically predict functional genes^19^. Here, we established a supervised ensemble machine learning Matlab package called FuncProFinder to predict meiosis-essential candidates based on the meiotic proteomic data (Methods and Supplementary Notes). According to phenotype annotation in MGI database, a protein is determined to be meiosis-essential if knockout of the protein leads to meiosis arrest, while a non-essential protein is termed that the protein knockout mice does not have lethal or meiosis-arrest phenotype. From the 8,002 identified proteins in this study, a total of 159 proteins were essential and 2,151 were non-essential (Fig. 3a; Supplementary Information, Table S3).

**Fig. 3.**
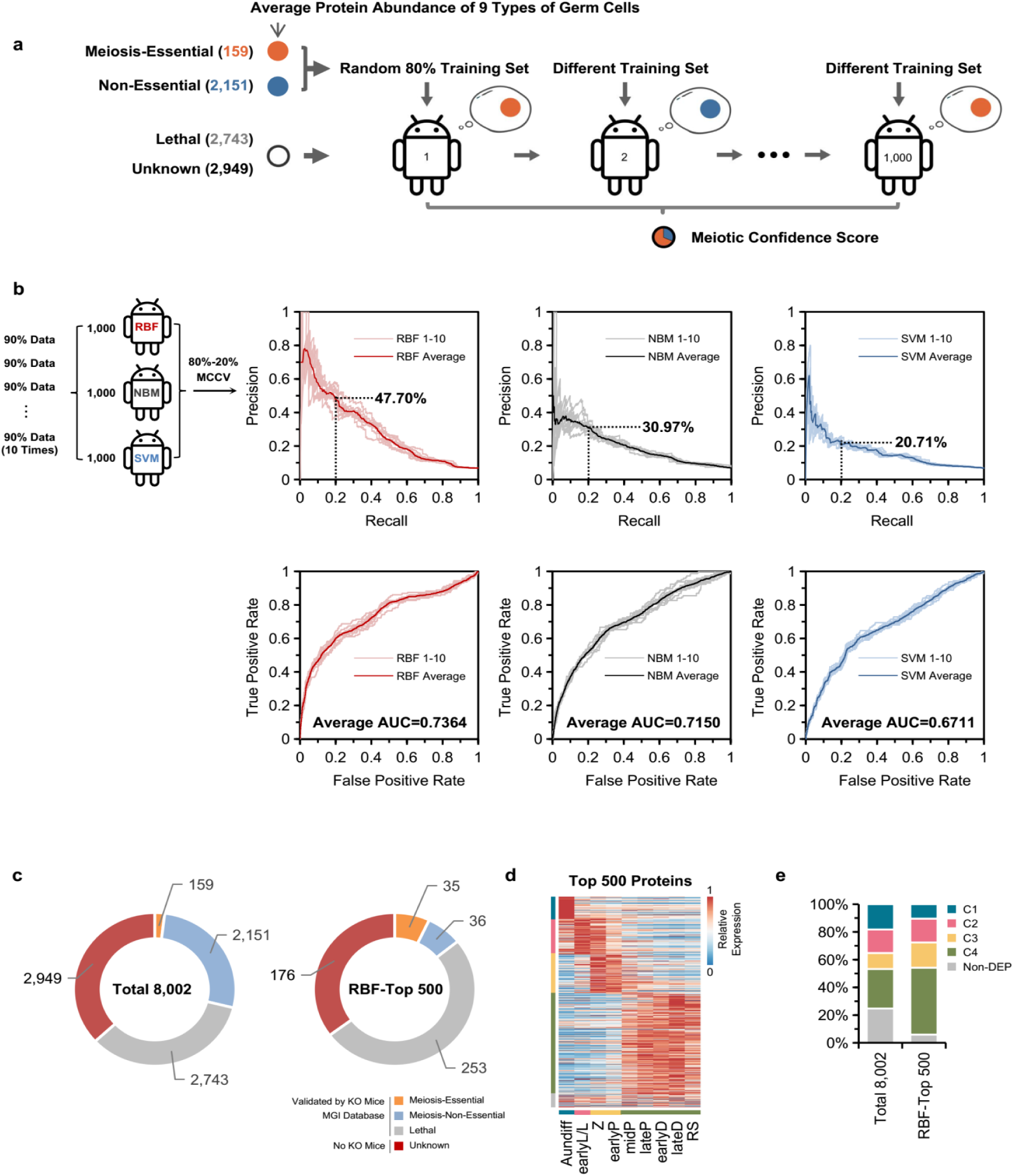
Prediction of meiotic-essential candidates using machine learning. **a** Workflow to build machine learning algorithms based on proteome data. **b** Comparison of the prediction results, Precision-Recall curve (upper panels) and ROC curve (lower panels) generated from three sub-classifiers, regularized RBF, NBM and SVM. MCCV, the Monte-Carlo cross validation. **c** On the basis of annotation of MGI database, the distribution of meiosis-essential, meiosis-non-essential, lethal and unknown proteins in 8,002 quantified proteins (left panel) and the top 500 meiosis candidates derived from RBF prediction (right panel). **d** Heatmap of dynamic abundance of the top 500 meiosis candidates derived from RBF prediction. **e** Comparison of the relative distribution of the DEP clusters treated with/without RBF filtration.

With abundances of these proteins as training sets, three methods of the FuncProFinder, radial basis function (RBF)^29^, naive Bayesian model (NBM), and support vector machine (SVM)^30^, were used to construct classifiers to predict whether or not a given protein was meiosis-essential. Based on the FuncProFinder package, the prediction precision tested by Monte-Carlo cross validation^31^ reached 47.70% (RBF), 30.97% (NBM), and 20.71% (SVM) with the recall setting at 0.2. AUCs (areas under curve) for the receiver operation characteristic (ROC) curve of the predictions were 0.7364 (RBF), 0.7150 (NBM), and 0.6711 (SVM) (Fig. 3b). As the prediction performance of RBF in both precision and AUC was superior to the other two algorithms in this dataset, FuncProFinder-RBF was accepted to predict the meiosis-essential possibility of a given protein.

Next, the identified proteins were scored with the FuncProFinder-RBF algorithm, and the higher the meiotic confidence scores, the more likely a protein was meiosis-essential (Supplementary Information, Table S3). A total of 500 proteins with the top scores were filtered and their functional information in MGI are shown in Fig. 3c. The ratio of meiosis-essential proteins against both essential and non-essential proteins was 6.54% (159:2310) for the all 8,002 proteins, whereas the ratio changed to 49.30% (35:71) for the top 500 selected candidates, indicating that meiosis-essential proteins could be greatly enriched. The top 500 candidates exhibited dynamic protein abundance changes during spermatogenesis (Fig. 3d), containing more DEPs (94.00%) compared to the total 8,002 proteins (75.23%). Additionally, in the top 500 candidates, the ratio of genes showed higher expression in the three meiotic cluster (C2-C4) was also higher than the ratio in the global proteome (Fig. 3e), consistent with the hypothesis that a meiosis-essential candidate could be a DEP with higher expression abundance in sub-phases of meiosis. Based on these results, the RBF algorithm is a potential method to select meiosis-dependent candidates from a large pool of identified proteins.

### Pyruvate dehydrogenase alpha 2 (*PDHA2*) is essential for meiosis

Of the top 500 candidates, a total of 176 proteins have never been knocked out for phenotype examination using mouse model according to the MGI database (Fig. 4a). Understanding of the functional pathways enriched in those predicted meiosis-essential candidates could assist in understanding the molecular basis in meiosis. Enrichr^32^, a pathway enrichment tool, was implemented to identify enriched pathways of these 176 candidates based on a KEGG 2018 mouse database. The top six functional categories after enrichment analysis were determined, and only one common metabolism pathway, pyruvate metabolism, was highly enriched in these meiosis-essential candidates (Fig. 4b). It has been reported that pyruvate metabolism was required in the isolated pachytene spermatocytes cultured *in vitro*^33^. However, whether pyruvate-related proteins were essential in meiosis has not been verified *in vivo*.

**Fig. 4.**
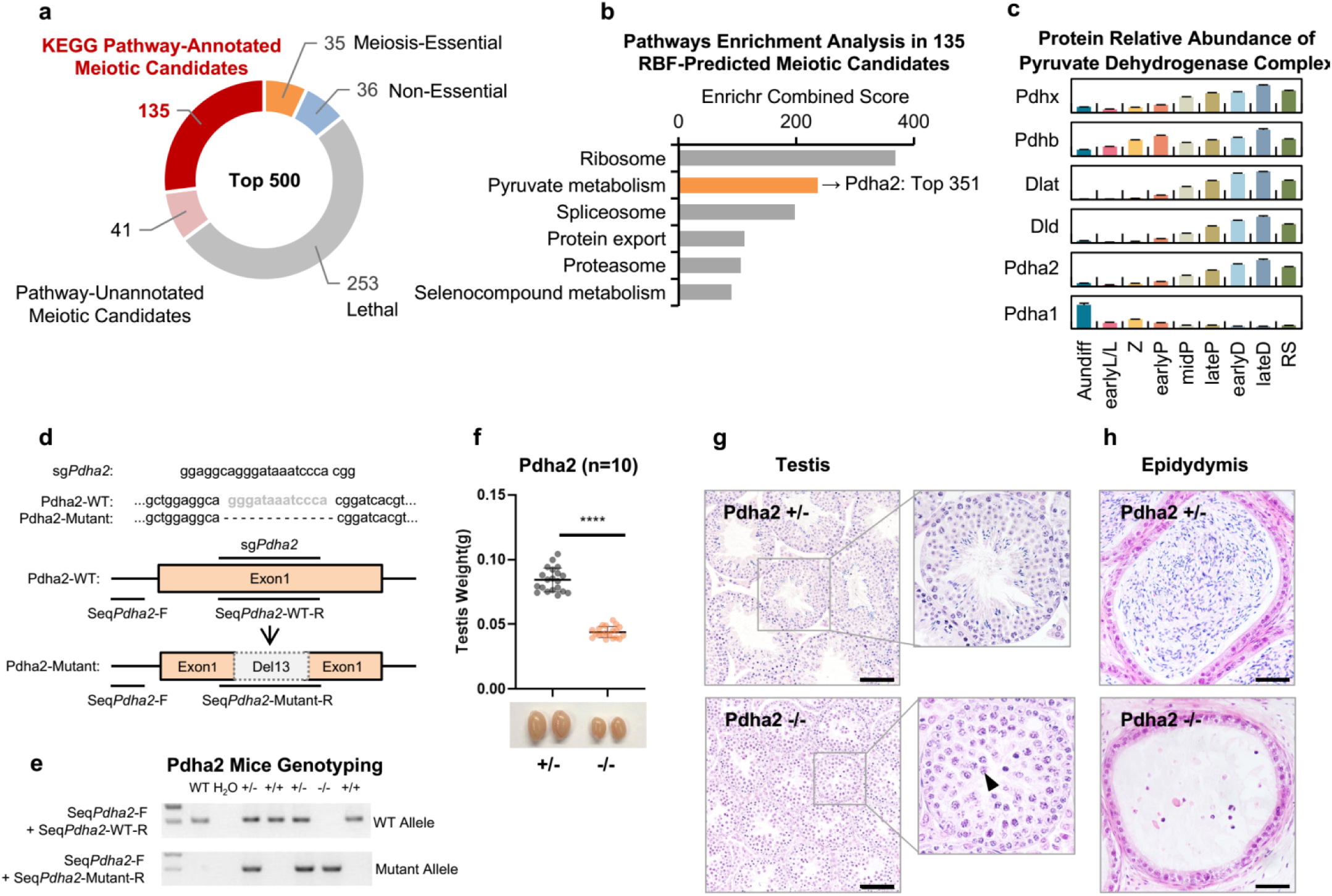
Phenotypic validation of the PDHA2 during meiosis. **a** Proteins with clearly functional annotation of the 176 proteins without KO evidence in the MGI database. **b** The KEGG pathways enriched in the proteins indicated in Fig. 4a. **c** The dynamically relative abundance of the six proteins of the PDC complex. The error bar represents SEM in from triplicate samples. **d** The strategy to construct Pdha2 knockout mouse. **e** Genotyping of Pdha2 knockout mice verified by PCR. **f** Comparison of the testes weights between Pdha2^+/-^ and Pdha2^-/-^ mice (n=10, unpaired two-tailed t test, p< 0.0001). **g** Cross-sections of H&E stained seminiferous tubules from 8-week-old Pdha2^+/-^ (upper panel) and Pdha2^-/-^ mice (lower panel), insets denote the specific seminiferous tubule under higher magnification. The arrow points to a Pachytene-like cell. **h** Cross-sections of epididymis from 8-week-old Pdha2^+/-^(upper panel) and Pdha2^-/-^ (lower panel) mice stained with H&E. Scale bar = 50 µm.

*PDHA2*, among the top-500 high-ranked meiosis-essential candidates, is a catalytic subunit of the pyruvate dehydrogenase complex (PDC), associated with four other proteins, *PDHB, DLAT, DLD*, and *PDHX*^34^. Lack of any component in the complex could lead to activity loss. In a previous study, the transcription status of the *Pdha2* gene was shown dynamically changed during spermatogenesis, increasing from the pachytene and gradually decreasing in spermatids^35^. With our proteomic data, the expression changes of all proteins in the PDC during spermatogenesis are further illustrated (Fig. 4c). The protein abundance of two catalytic subunits of PDC – *PDHA1* and *PDHA2* – changed in opposite directions: X-chromosome-linked protein *PDHA1* decreased during meiosis due to meiotic sex chromosome inactivation (MSCI), whereas *PDHA2* increased from earlyP to lateD. For the other four components of PDC, the changes in their abundance were similar to *PDHA2*, implying that the PDC had structural integrity of catalytic functions during meiotic development.

To further verify the physiological roles of *Pdha2* during meiosis, a *Pdha2* knockout mouse model was generated by CRISPR/Cas-mediated genome engineering. A 13-basepair deletion was induced into the *Pdha2* exon, which led to a reading-frame shift of *Pdha*2 and early termination (Fig. 4d); the knockout result was examined by genotyping (Fig. 4e). The testes weights of 8-week-old adult *Pdha2*^-/-^ mice were significantly smaller than *Pdha2*^+/-^ mice (Fig. 4f), and the testes histology of these mice was further examined by H&E staining of cross sections (Fig. 4g, h). In *Pdha2*^+/-^ mice, the testes were comparable to WT mice, in which all types of germ cells were observed, and mature sperm had fully filled their epididymides. In contrast to their heterozygous littermates, *Pdha2*^-/-^ mice entirely lacked post-meiotic cells, and pachytene-like spermatocytes were accumulated in their testes. Furthermore, no spermatozoa were observed in their epididymides. Thus, with the help of FuncProFinder-RBF prediction, this study provides evidence that *PDHA2* is a meiotic- regulation factor, knockout of which is likely to stop the meiosis process at the pachytene stage. As PDC catalyzes pyruvate to acetyl-CoA and determines the energy level in a cell, it is a reasonable deduction that *PDHA2*, as a key component of PDC, could regulate ATP generation in spermatocytes and affect the meiotic process.

### Phenotype verification of the top ten male meiosis-essential candidates without functional annotation

Among the 176 proteins mentioned above, 41 proteins appeared to lack KEGG pathway annotation (Fig. 5a). Whether they are essential for meiosis needs to be further verified by experiments. The top ten candidates based on meiotic confidence score were selected and knocked-out in mice using CRISPR/Cas-mediated genome engineering (Supplementary Information, Fig. S3a-j). Abundance changes of the ten proteins are shown in Fig. 5b. After knock-out treatment, we obtained surviving homozygous deficient pups from eight of the ten genes, as the deficiency of the *Gapvd1* and *1700037H04Rik* in mice were lethal. The reproductive anatomy of these non-lethal mice were carefully examined by testis weight and histological analysis. In the *Txnl1*^-/-^, *AA467197*^-/-^, *Lrrc40*^-/-^, and *Naxe*^-/-^mice, no significant change was observed in their testes weights or histology (Supplementary Information, Fig. S3a-h). However, knockout of the other four genes affected the meiosis process of the homozygous deficient mice. Generally, the testis weights of the *Zcwpw1*^-/-^, *Tesmin*^-/-^, *Kctd19*^-/-^, and *1700102P08Rik*^-/-^ mice were significantly lighter than their heterozygous littermates (Fig. 5c-f). Specifically, H&E stained images of the testes derived from *Zcwpw1*^-/-^, *Tesmin*^-/-^, and *1700102P08Rik*^-/-^ mice appeared to have a pachytene-arrested phenotype, pachytene spermatocytes with condensed nuclei, lack of post-meiotic cells, and tubules that were highly vacuolized (Fig. 5g-i). The *Kctd19*^-/-^ mice exhibited a typical metaphase I–arrested phenotype, containing spermatocytes from leptotene to metaphase I, but with no post-meiotic cells (Fig. 5j).

**Fig. 5.**
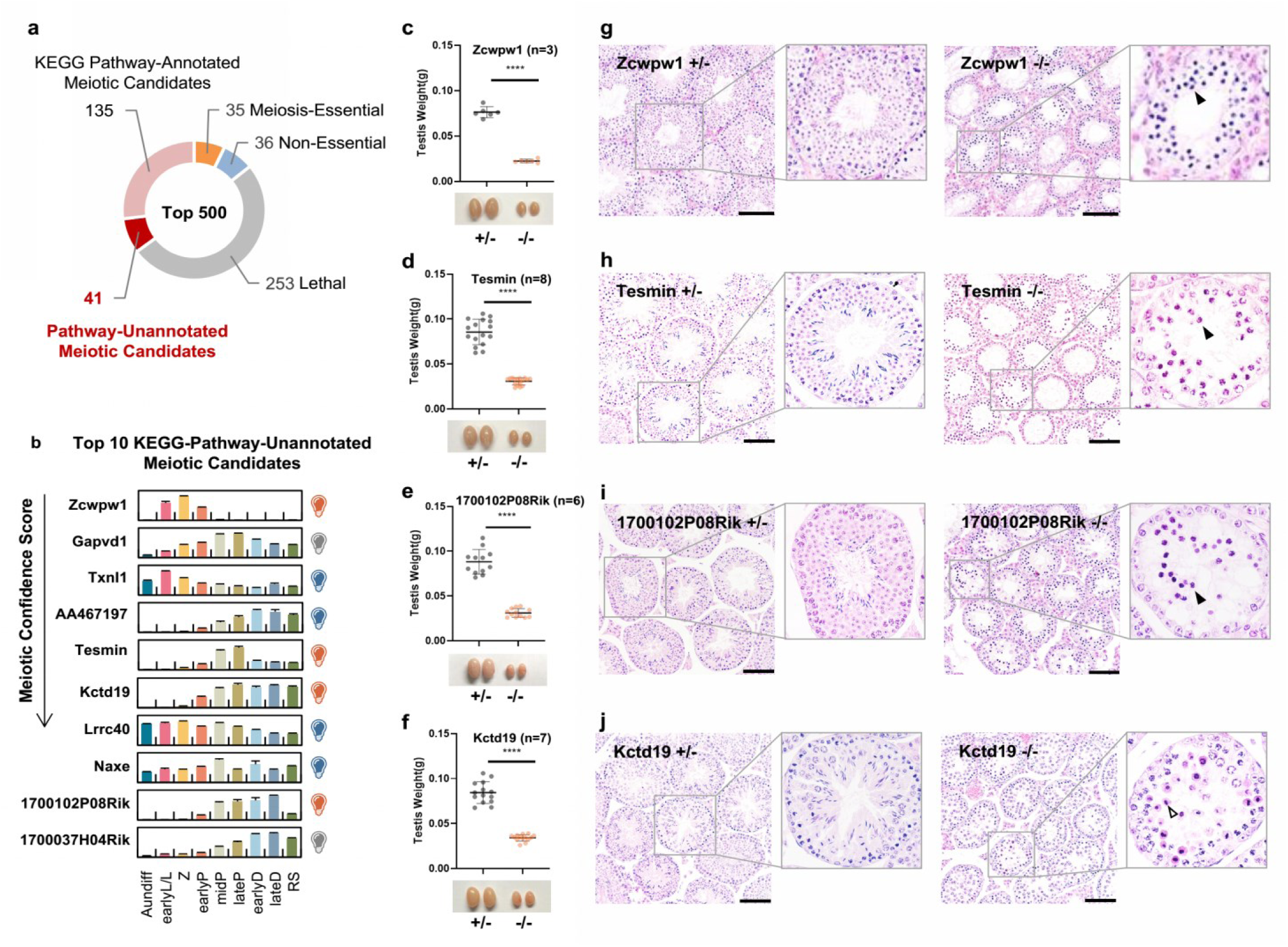
Phenotypic validation of the meiotic-essential proteins predicted by FuncProFinder-RBF. **a** Proteins without functional annotation of the 176 proteins without KO evidence in the MGI database. **b** The dynamic relative abundance of the top 10 RBF-ranked candidates during spermatogenesis. The error bar represents SEM in triplicates. The bulbs on the figure (right) indicate the phenotype of KO mice, orange as meiosis-essential, blue as meiosis non-essential and grey as lethal. **c-f** Comparison of testis weights derived from 8-week-old Zcwpw1+/- and Zcwpw1-/- mice (**c**), Tesmin+/- and Tesmin -/- mice (**d**), Kctd19+/- and Kctd19-/- mice (**e**), and 1700102P08Rik+/- and 1700102P08Rik -/- mice (**f**). **** represents p< 0.0001 in unpaired two-tailed *t*-test. **g-j**. Cross-sections of H&E stained seminiferous tubules from the heterozygous and homozygous knockout mice of the four genes above, insets denote the specific seminiferous tubule under higher magnification. The filled arrow points to a Pachytene-like cell. The hollow arrow points to a Metaphase I-like cell. Scale bar = 50 µm.

The molecular mechanism of pachytene arrest is hypothesized to result from failure of DNA repair or incomplete synapsis^36, 37^. As knock out of *Zcwpw1, Tesmin*, or *1700102P08Rik* led to pachytene arrest, the molecular mechanisms underlying the pachytene arrest phenotype in these three knockout mice lines need to be verified. To address this question, the spermatocytes in *Zcwpw1*^-/-^, *Zcwpw1*^+/-^, *Tesmin*^-/-^, *Tesmin*^+/-^, *1700102P08Rik*^-/-^, or *1700102P08Rik*^+/-^ mice were chromosome-spread and immunostained with antibodies against DNA repair and synapsis events, including *SYCP3* and *SYCP1* as components of the SC, *γH2AX* as an indicator of DNA damage, and *MLH1* as a marker of crossover formation. The immunostaining of the four different antibodies against the spermatocytes from *Tesmin* and *1700102P08Rik* in both heterozygous and homozygous knock out mice exhibited no difference (Supplementary Information, Fig. S4f-n), implying that neither DNA repair nor synapsis were affected by gene knockout. However, the immunostaining of these antibodies against the *Zcwpw1*^-/-^ spermatocytes was quite different from *Zcwpw1*^+/-^ (Fig. 6a-e). The staining signal distribution of *γH2AX* in the leptotene spermatocytes of the *Zcwpw1*^-/-^ mice was comparable with that of *Zcwpw1*^+/-^, suggesting that the formation of double strand breaks (DSBs) was not affected by the absence of *ZCWPW1* (Fig. 6a). In the pachytene spermatocytes, the *γH2AX* staining was only seen in the sex body region, whereas it was still spread out in the autosome regions in *Zcwpw*1^-/-^, indicating that the DSB repair was not finished on the autosome without *ZCWPW1* (Fig. 6b). Co-immunostaining of *SYCP3* and *SYCP1* were highly merged in the chromosomes in *Zcwpw1*^+/-^ mice, whereas the locations of the two SC components were not fully overlapped in *Zcwpw1*^-/-^ mice, suggesting that the SC were not fully formed due to the lack of *ZCWPW1* (Fig. 6c). Furthermore, the *MLH1* loci were perceived in *Zcwpw1*^+/-^ pachytene spermatocytes and were counted within a normal range, whereas the *Zcwpw1*^-/-^ pachytene spermatocytes appeared to have nearly no *MLH1* loci, implicating that crossovers were not formed without *ZCWPW1* (Fig. 6d, e). Based on these results, *ZCWPW1* functions directly in both DNA repair and synapsis, but the functions of *1700102P08Rik* and *Tesmin* are not clearly clarified even though the two proteins participate in the regulation of meiosis.

**Fig. 6.**
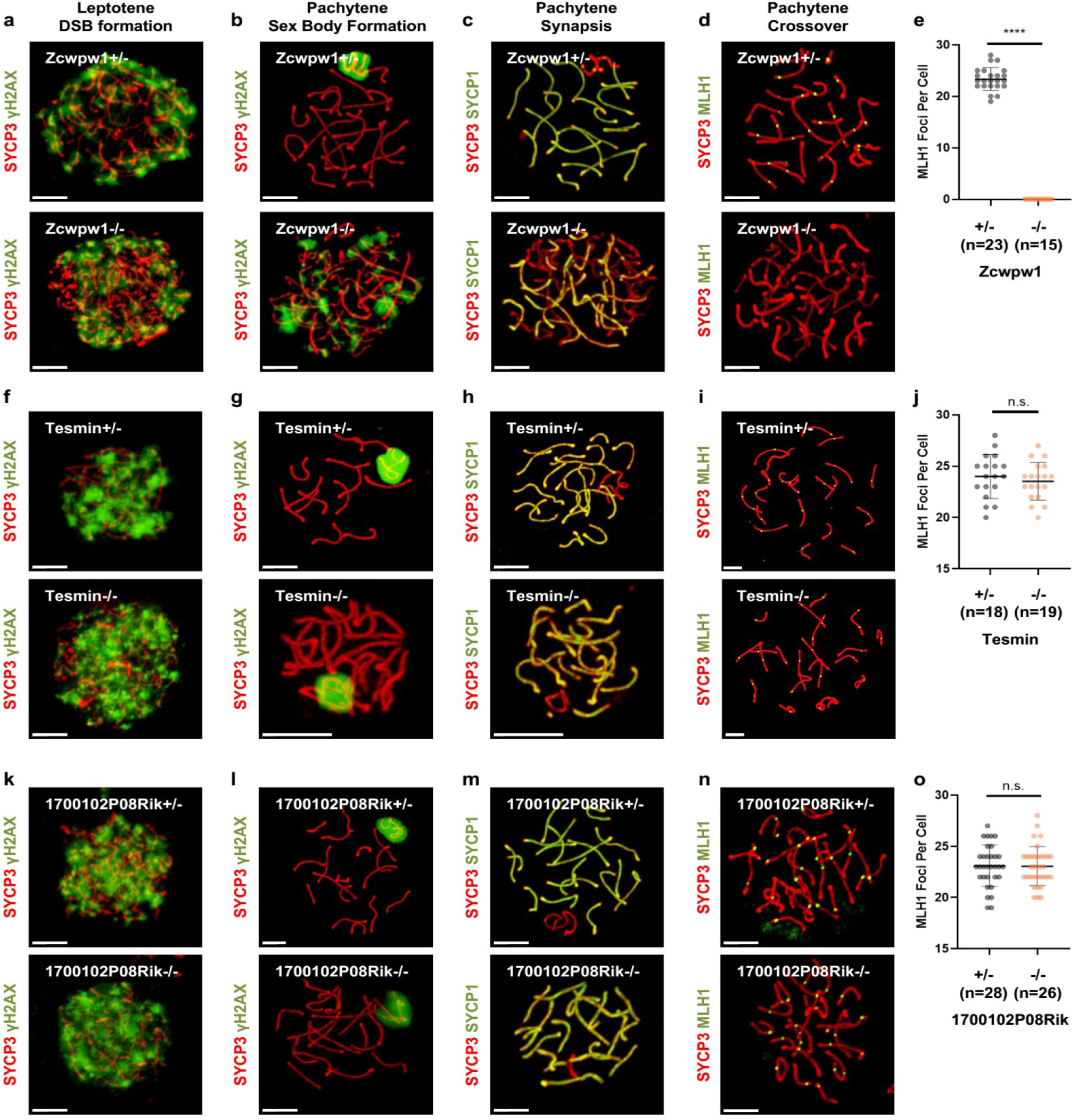
Analysis of the homologous recombination and synapsis states by confocal images in the heterozygous and homozygous gene knockout mice, Zcwpw1 (a-d), Tesmin (f-i) and 1700102P08Rik (k-n). Confocal images in the two columns on the left side, the cells were co-immuno-stained with anti-SYCP3 (red) and anti-γH2AX (green); in the next column with anti-SYCP3 (red) and anti-SYCP1 (green), and in the furthest right column with anti-SYCP3 (red) and anti-MLH1 (green). Scale bar = 50 µm. **e-o** Comparison of MLH1 foci number in spermatocytes derived from Zcwpw1^+/+^ and Zcwpw1^−/−^ (**e**), Tesmin ^-/-^ and Tesmin ^+/-^(**j**), 1700102P08Rik ^+/-^ and 1700102P08Rik ^-/-^ (**o**) mice. **** represents p< 0.0001 in unpaired two-tailed *t*-test; n.s. means no significance.

To summarize the knockout experiments for predicting the meiosis-essential proteins, FuncProFinder-RBF offered a set of satisfactory candidates. Of the ten candidates, at least 40% were verified as meiosis-essential. As their functions are not annotated yet, their involvement in meiosis would be an interesting direction for functional exploration. For example, the pachytene arrest in *Zcwpw*1^-/-^ mice was found to result from failed DSB repair and incomplete synapsis.

## DISSUCSSION

In this study, one of the fundamental goals was to acquire global and quantitative proteomic profiling during different stages of mouse meiosis. How to obtain such information is a long-lasting question. Proteomic investigations of the entire mouse testis and one type of meiotic cells – pachytene spermatocytes – have been accomplished in several labs^7, 18, 38^. However, without isolation of different types of meiotic cells, these studies could not generate a precise picture of the different stages of mouse meiosis.

Here, first, we designed a project that enabled comprehensive proteomic profiling around meiosis. To reach our goal, seven consecutive types of meiotic cells plus pre-meiotic spermatogonia and post-meiotic round spermatids were isolated and proteins in each cell type were identified and quantified by high-resolution mass spectrometry. A total of 8,002 proteins were identified, including 6,020 differentially expressed proteins, which is the largest dataset of proteomics related to meiosis, offering global information of protein quantities in different stages of the mammalian meiosis process.

Second, this comprehensive proteomics is likely to provide new views for understanding of meiosis. For instance, it is generally accepted that spermatocytes are categorized in several sub-stages judged by the status of chromosome morphology^23, 39^. Based on this criterium, earlyP and midP are categorized to the similar group of pachytene cells; nevertheless, the protein abundance correlation revealed that these two types of spermatocytes with similar appearance were totally different (Fig. 2a, d). For another example, components of the SC were assumed to mainly exist from Z to lateP and decreased after lateP. However, *SYCE1*, a key SC component, was consistently detected after lateP to RS with relatively high abundance (Fig. 2c), implying that *SYCE1* might perform additional functions (except synapsis) after meiosis Prophase I. Therefore, the refined profile of quantitative proteome provided a new assessment of molecular events related to meiosis.

Third, informatics analysis of proteomic data related to spermatogenesis is likely to provide a basis for functional exploration of mouse meiotic genes. Genes involved in critical meiotic events, such as homologous recombination and synapsis, are not well indentified^1, 2^. Miyata et al. selected 54 testis-specific genes and constructed the correspondent gene knockout mice, but unfortunately, they did not find any gene essential to meiosis^8^. In this study, an ensemble strategy of machine learning was used to predict meiosis-essential proteins based on the different abundance patterns between meiosis-essential or non-essential proteins. With this strategy, enrichment of meiosis-essential proteins was raised from 6.54% to 49.30% after prediction on the test set (Fig. 3c). Moreover, at least four of top ten candidates without KEGG pathway annotation were confirmed as meiosis-essential proteins by gene-knockout mouse (Fig. 4c-j). Hence, functional exploration based upon proteomics seems to significantly improve prediction efficiency.

In addition, with the machine learning prediction and gene knockout validation, meiosis development of three genes, *Zcwpw1, Tesmin* and *1700102P08Rik*, was found to be arrested at pachytene in homologous gene knockout mice. Immuno-fluorescence images in this study revealed that *Zcwpw1* functioned in DNA repair and synapsis. Very recently, *Zcwpw1* was identified as a histone H3K4me3 reader required for repair of PRDM9-dependent DNA double strand breaks and synapsis by three independent research groups^40-43^, which was consistent with our observations. Knockout of the other two genes, *Tesmin* and *1700102P08Rik*, were found to have no effect on DNA repair and synapsis, indicating they could be involved in presently unknown molecular events inspected by midPachytene checkpoint. These meiosis-essential biological events need further exploration. Additionally, a list of meiosis-essential candidates without KEGG annotation were presented here (Supplementary Information, Table S3). Based on this list, more meiosis-essential genes could be verified, and new biological events could be uncovered concerning the molecular basis of mouse meiosis.

## METHODS

### Mice for experiments

Wild-type mice, C57BL/6Slac, were purchased from SLAC China. All animal experiments were conducted in accordance with the guidelines of the Animal Care and Use Committee at Shanghai Institute of Biochemistry and Cell Biology, Chinese Academy of Science.

### Spermatogenesis synchronization

Spermatogenesis was synchronized as previously described with modifications^9, 21, 22^. Briefly, C57BL/6Slac mice, from P2 to P8, were fed on WIN 18,446 suspended in 1% gum tragacanth at 100 μg/g body weight, which blocks spermatogonia differentiation and synchronizes the spermatogenesis process. Spermatogonia differentiation was re-initiated in these mice at P9 through intraperitoneal injection of retinoic acid in dimethyl sulfoxide at 35 μg/g body weight. The testes of P37 to P46 mice were collected and evaluated for efficiency of synchronization with histological analysis and cell sorting.

### Isolation of mouse spermatocytes

Spermatocytes were isolated by fluorescent-activated cell sorting (FACS) as previously described with modifications^23^. Briefly, testes of an individual spermatogenesis-synchronized mouse were collected in GBSS. After removal of the tunica albuginea, the testes were incubated in 5 ml of DMEM containing 120 U/ml of collagenase type I at 32 °C with gentle agitation for 15 min. The dispersed seminiferous tubules were further digested with 5 ml of 0.15% trypsin and DNase I (10 μg/ml) at 32 °C for 30 min, and the digestion was terminated by adding 0.5 ml of fetal bovine serum (FBS). The suspension of dissociated testicular cells was filtered through a DMEM-prewetted cellular filter, followed by centrifugation at 500 ×g for 5 min at 4 °C. The cells were resuspended in DMEM with Hoechst 33,342 (5 mg/ml) to concentrations of 1 × 10^6^ cells/ml, and were treated with propidium iodide (2 mg/ml) and DNase I (10 μg/mL) in a rotator for 30 min at 32 °C at 10 rpm/min. The treated cells were centrifuged at 500 ×g for 5 min at 4 °C and resuspended in 1 ml DMEM for sorting based on their Hoechst 33,342 staining using a FACSAria II cell sorter (BD Biosciences).

### Isolation of mouse THY1+ c-KIT-spermatogonia

Undifferentiated spermatogonia (THY1+ c-KIT-spermatogonia) were isolated using magnetic activated cell sorting (MACS) with magnetic microbeads conjugated to anti-THY1 (130-049-101, Miltenyi Biotech) and anti-c-KIT (130-091-224, Miltenyi Biotech) as described previously^20^. Briefly, testes of P7 mice were digested with collagenase type I and trypsin. After digestion, the testis cells were suspended in PBS, then layered on 2 ml 30% Percoll, followed by centrifugation at 600 ×g for 8 min. The cell pellets were resuspended with PBS containing anti-c-KIT magnetic microbeads. With 20 min incubation, the mixtures were loaded on a magnetic device to collect c-KIT-cells. These c-KIT-cells were incubated with anti-THY1 magnetic microbeads for 20 minutes, and the THY1+ c-KIT-cells were enriched using MS columns (130-042-201, Miltenyi Biotech) and a MiniMACS separator (130-042-102, Miltenyi Biotech). The purity of THY1+ c-KIT-cells were estimated by anti-PLZF (SC-22839, Santa Cruz Biotechnology) and DAPI staining.

### Histological analysis

Testes were fixed in Bouin’s solution, embedded in paraffin and sectioned. The sections were dewaxed with xylene and were re-hydrated in a series of ethanol concentrations. The treated sections were stained with hematoxylin and eosin (H&E) and were sealed with nail polish. Spermatogenesis stages in seminiferous tubule cross-sections were recognized as previously described^39^.

### Meiotic chromosome spreading and immunofluorescence

Meiotic spreads were made following a previously described protocol with modifications^44^. Briefly, the cells were re-suspended in hypotonic extraction buffer, then in 100 mM sucrose. The cell suspension was pipetted onto glass slides that were coated in thin layer of 1% PFA and 0.15% Triton X-100. The slides were dried slowly in a humid chamber at room temperature. For immunofluorescence staining, the slides were washed with PBS and blocked with Tris-HCl buffer saline containing 0.5% Tween-20 and 3% BSA for 30 min, and were incubated with different antibodies, as anti-SCP1 (ab15090, Abcam), anti-SCP3 (sc-74569, Santa Cruz), anti-SCP3 (ab15093, Abcam), anti-γH2AX (05-636, Millipore), anti-γH2AX (#9718, CST) and anti-MLH1 (550838, BD Pharmingen). Finally, the slides were incubated with Alexa Fluor 488- or 594-conjugated secondary antibodies (711545152 and 711585152, Jackson ImmunoResearch Laboratories) to detect meiotic-relevance signals, and were treated with DAPI for defining nuclei. The spread cells were monitored under confocal laser scanning microscope FV3000 (Olympus).

### Protein extraction and digestion

The cells were homogenized by pipetting in lysis buffer containing 7 M urea, 2 M thiourea, 0.2% SDS, 100 mM Tris-HCl, 10 mM DTT and 1x cocktail-free protease inhibitor (Promega), pH 7.4. The proteins in lysates were reduced with 5 mM DTT and were alkylated with 55 mM IAM, then were further extracted by cold acetone precipitation. The precipitated proteins were resolved in 7 M urea lysis buffer, and the protein concentrations were estimated using a Bradford protein assay (Bio-Rad). For each sample, 200 μg protein was loaded into 10 kD spin filters (Millipore) and centrifuged at 12,000 g for 20 min. The filter was washed in order with urea lysis buffer and 1 M TEAB and then treated with the trypsin (Promega) at a ratio of 1:50 at 37°C for 16 hours, shaking at 600 rpm in a Thermomixer (Eppendorf). Tryptic peptides were collected by centrifugation and were quantified using a Quantitative Colorimetric Peptide Assay (Thermo Scientific).

### Peptide fractionation on RP-HPLC

For each sample, approximately 100 μg peptides were dissolved in elution buffer A containing 20 mM ammonium bicarbonate and 5% acetonitrile, pH 9.8. The dissolved peptides were loaded on a Phenomenex C18 column (5 μm particle, 110 Å pore and 250 mm × 4.6 mm) that was mounted on a Shimadzu liquid chromatography system and pre-equilibrated with elution buffer B containing 20 mM ammonium bicarbonate and 90% acetonitrile, pH 9.8. The peptides were eluted through a stepped gradient program as follows: 0–3 min, 5% B; 3–7 min, 9% B; 7–11 min, 13% B; 11–15 min, 19% B; 15–19 min, 80% B; 19-21 min, 5% B; 21-21.5 min, 5%-80% B; 21.5-22.5 min, 80% B; 22.5-23 min, 80%-5% B and 23-29 min, 5% B at a flow of 1 ml/min. Twenty-four fractions from 3-26 min were collected and these fractions were further combined to five fractions according to the absorption peaks at 214 nm during chromatography, fractions 1-5 as F1, 6-10 as F2, 11-14 as F3, 15-18 as F4 and 19-24 as F5.

### Peptide detection by LC-MS/MS

Identification of peptides was conducted on a quadrupole Orbitrap mass spectrometer (Q Exactive HF, Thermo Fisher Scientific) coupled to an UHPLC system (Dionex Ultimate 3000, Dionex) via a nanospray Flex ion source. About 1 μg of peptides were loaded on an C18 trap column (75 μm I.D. × 1.5 cm; in-house packed using Welch C18 3 μm silica beads) and were directly loaded into an C18 analysis column (75 μm I.D. × 20 cm; in-house packed using Welch C18 3 μm silica beads). The peptides used for mass spectrometry were eluted at 300 nl/min and 40 °C with two elution buffers, buffer A: 0.1% formic acid and 2% acetonitrile, and buffer B: 0.1% formic acid and 98% acetonitrile, following a gradient program, 0–5 min, 5% B, 5–7 min, 5– 7% B, 7–67 min, 7-28% B, 67-80 min, 28-43% B, 80-82 min, 43%-98% B, 82-84 min, 98% B, 84-85 min, 5% B and 85-90 min, 5% B. The mass spectrometer was operated in “top-30” data-dependent mode, collecting MS spectra in the Orbitrap mass analyzer (120,000 resolution at 350-1500 *m/z* range) with an automatic gain control (AGC) target of 3E6 and a maximum ion injection time of 50 ms. The ions with higher intensities were isolated with an isolation width of 1.6 *m/z* and were fragmented through higher-energy collisional dissociation (HCD) with a normalized collision energy (NCE) of 28%. The MS/MS spectra were collected at 15,000 resolution with an AGC target of 1E5 and a maximum ion injection time of 45 ms. Precursor dynamic exclusion was enabled with a duration of 60 s.

### Peptide search and protein quantification by Maxquant

Tandem mass spectra were searched against the 2018 Swiss-Prot mouse databases (downloaded 11-19-2018) using MaxQuant (v 1.5.3.30)^45^ with a 1% FDR at peptide and protein level. The search parameters for a peptide were set as trypsin digestion only, maximum of two missed cleavages of trypsin, minimum length of six amino acids, cysteine carbamidomethylation as fixed modification, N-terminal acetylation and methionine oxidations as variable modifications. The “Match Between Run” option was used. Label-free quantification (LFQ) was estimated with MaxLFQ algorithm, using a minimum ratio count of 1, and the specifically relative LFQ for a protein was defined by the ratios of the protein LFQ at certain sub-stage being divided by the protein maximum LFQ.

### Proteomic informatics analysis

Bioinformatics analysis of the identified and quantified proteins was performed with Perseus software (version 1.6.1.3)^28^, R statistical software and excel. Differentially expressed proteins (DEPs) among sub-stages were defined by two-way ANOVA analysis in Perseus filtered with adjusted q value < 0.001. To look for the DEP groups with protein abundance changes during spermatogenesis that share similar patterns, the DEPs were first evaluated by NbClust package in R (version 3.5.1)^46^ to find the optimum K value, and the relative LFQ of DEPs were clustered using K-means analysis. Gene Ontology and KEGG pathway analysis were performed using David GOBP (version 6.8)^47^ and Enrichr^32^, in which an enriched function was accepted upon p values less than 10e-8 and only the GO or pathway terms with <250 gene number in that gene sets were included. Heatmap data was visualized using Seaborn 0.9.0^48^.

### Machine learning for prediction of meiosis-essential proteins

Three sub-classifiers, regularized RBF, NBM or SVM, were employed to gain better predictions for meiosis-essential protein candidates. Protein abundance in nine types of germ cells were inputted to the classifiers to train the weights. The meiosis-essential proteins, defined as the proteins that overlapped the MGI database and proteomics, in this study were labeled as positive and non-essential proteins were labeled as negative. The ensemble learning process was broadly conducted in two steps as described previously with modifications^49^, the details of regularized RBF, NBM and SVM are presented in Supplementary Notes:

Step 1. All labeled proteins were randomly divided into two sets, training (80%) and testing (20%). A single sub-classifier was constructed by the training set and the corresponding output function were generated for prediction. The predicted score for the proteins in the testing set were estimated by the output function. This process was repeated 1,000 times to construct the ensemble algorithm.

Step 2. To test the performance of the algorithm, a Monte-Carlo cross validation was applied^31^. The final predicted score of each protein, called the meiotic confidence score, is defined as the mean value of the individual scores in step 1. Precision-recall curve and receiver operating characteristic (ROC) curve were used to evaluate the prediction performance of the three algorithms based on regularized RBF, NBM and SVM.

The FuncProFinder Matlab package was uploaded on: https://github.com/sjq111/FuncProFinder.

### Generation of gene knockout mice with the CRISPR/Cas9 system

For generation of knockout mice corresponding to the genes of interest in this study, sgRNAs were designed based on their genome structures (listed in Supplementary Information, Table S4). T7-Cas9 PCR product was gel purified and used as the template for *in vitro* transcription (IVT) using mMESSAGE mMACHINE™ T7 ULTRA transcription kit (AM1345, ThermoFisher Scientific). The T7-sgRNA PCR product was gel purified and used as the template for IVT with a MEGAshortscript™ T7 transcription kit (AM1354, ThermoFisher Scientific). Both the Cas9 mRNA and the sgRNAs were purified using MEGAclear™ Transcription Clean-Up Kit (AM1908, ThermoFisher Scientific) and eluted in RNase-free water (10977015, ThermoFisher Scientific). The Cas9 mRNA and sgRNAs were injected into one-cell embryos as described previously^50-52^. The injected embryos were cultured *in vitro* to develop to 2-cell embryos and transplanted to oviducts to generate knockout pups. After pups were born, genotyping was performed by direct sequencing following PCR to validate the knockout consequences. Genotyping primer sequences that were used are listed in Table S4.

### Data availability

The mass spectrometry proteomics data have been deposited to the ProteomeXchange Consortium via the PRIDE^53^ partner repository with the dataset identifier PXD017284.

## ACKNOWLEDGEMENTS

We thank Dr. Dangsheng Li (Shanghai Institute of Biochemistry and Cell Biology) for helpful discussion of the manuscript. Charlie Degui Chen was supported by the Strategic Priority Research Program of the Chinese Academy of Sciences (XDB19000000), and the National Natural Science Foundation of China (91753128, 81772472). Liu Siqi was supported by the National Key R&D Program of China (2017YFC0908400 and 2017YFC0906703), the National Natural Science Foundation of China (NO.31700728) and the Shenzhen Engineering Laboratory for Proteomics (DRC-SZ[2016]749). Yang Hui was supported by the R&D Program of China (2018YFC2000100 and 2017YFC1001302), CAS Strategic Priority Research Program (XDB32060000), National Natural Science Foundation of China (31871502, 31522037), Shanghai Municipal Science and Technology Major Project (2018SHZDZX05), and Shanghai City Committee of science and technology project (18411953700, 18JC1410100). We thank the Histology, Flow Cytometry, and Animal services at SIBCB.

## AUTHOR CONTRIBUTIONS

D.C. and K.F conceived the project. K.F performed the synchronization of mouse spermatogenesis, isolated different types of germ cells, validated the phenotype of knockout mice and contributed to the manuscript; W.G performed the experiments to produce the proteomic data and contributed to proteomic data analysis; Q.L performed the proteomic bioinformatic data analysis and wrote the manuscript; J.S developed the machine learning package FuncProFinder; Y.W, C.Z, R.W and W.Y constructed all the knockout mouse models. L.Y and Y.Z helped to the visualization of proteomic data. H.Y supervised the knockout mouse production; S.L supervised the proteomic data production, data analysis and refined the manuscript; D.C supervised the project. All authors contributed to the manuscript.

## REFERENCES

1. Cahoon, C.K. & Hawley, R.S. Regulating the construction and demolition of the synaptonemal complex. Nat Struct Mol Biol 23, 369–377 (2016).

2. Keeney, S., Lange, J. & Mohibullah, N. Self-organization of meiotic recombination initiation: general principles and molecular pathways. Annu Rev Genet 48, 187–214 (2014).

3. Romanienko, P.J. & Camerini-Otero, R.D. The mouse Spo11 gene is required for meiotic chromosome synapsis. Mol Cell 6, 975–987 (2000).

4. Baudat, F., Manova, K., Yuen, J.P., Jasin, M. & Keeney, S. Chromosome synapsis defects and sexually dimorphic meiotic progression in mice lacking Spo11. Mol Cell 6, 989–998 (2000).

5. Bannister, L.A. et al. A dominant, recombination-defective allele of Dmc1 causing male-specific sterility. PLoS Biol 5, e105 (2007).

6. Petukhova, G.V., Romanienko, P.J. & Camerini-Otero, R.D. The Hop2 protein has a direct role in promoting interhomolog interactions during mouse meiosis. Dev Cell 5, 927–936 (2003).

7. Wang, L. et al. Proteomic Analysis of Pachytene Spermatocytes of Sterile Hybrid Male Mice. Biol Reprod 95, 52 (2016).

8. Miyata, H. et al. Genome engineering uncovers 54 evolutionarily conserved and testis-enriched genes that are not required for male fertility in mice. Proc Natl Acad Sci U S A 113, 7704–7710 (2016).

9. Chen, Y. et al. Single-cell RNA-seq uncovers dynamic processes and critical regulators in mouse spermatogenesis. Cell Res 28, 879–896 (2018).

10. Green, C.D. et al. A Comprehensive Roadmap of Murine Spermatogenesis Defined by Single-Cell RNA-Seq. Dev Cell 46, 651–667 e610 (2018).

11. Lukassen, S., Bosch, E., Ekici, A.B. & Winterpacht, A. Characterization of germ cell differentiation in the male mouse through single-cell RNA sequencing. Scientific reports 8, 6521 (2018).

12. Jung, M. et al. Unified single-cell analysis of testis gene regulation and pathology in five mouse strains. Elife 8 (2019).

13. Ernst, C., Eling, N., Martinez-Jimenez, C.P., Marioni, J.C. & Odom, D.T. Staged developmental mapping and X chromosome transcriptional dynamics during mouse spermatogenesis. Nat Commun 10, 1251 (2019).

14. Guo, J. et al. The adult human testis transcriptional cell atlas. Cell Res 28, 1141–1157 (2018).

15. Hermann, B.P. et al. The Mammalian Spermatogenesis Single-Cell Transcriptome, from Spermatogonial Stem Cells to Spermatids. Cell Rep 25, 1650–1667 e1658 (2018).

16. Xia, B. et al. Widespread Transcriptional Scanning in the Testis Modulates Gene Evolution Rates. Cell 180, 248–262 e221 (2020).

17. Cagney, G. et al. Human tissue profiling with multidimensional protein identification technology. J Proteome Res 4, 1757–1767 (2005).

18. Gan, H. et al. Integrative proteomic and transcriptomic analyses reveal multiple post-transcriptional regulatory mechanisms of mouse spermatogenesis. Mol Cell Proteomics 12, 1144–1157 (2013).

19. Krishnan, A. et al. Genome-wide prediction and functional characterization of the genetic basis of autism spectrum disorder. Nat Neurosci 19, 1454–1462 (2016).

20. Kubota, H., Avarbock, M.R. & Brinster, R.L. Growth factors essential for self-renewal and expansion of mouse spermatogonial stem cells. Proc Natl Acad Sci U S A 101, 16489–16494 (2004).

21. Hogarth, C.A. et al. Turning a spermatogenic wave into a tsunami: synchronizing murine spermatogenesis using WIN 18,446. Biol Reprod 88, 40 (2013).

22. Romer, K.A., de Rooij, D.G., Kojima, M.L. & Page, D.C. Isolating mitotic and meiotic germ cells from male mice by developmental synchronization, staging, and sorting. Dev Biol 443, 19–34 (2018).

23. Gaysinskaya, V., Soh, I.Y., van der Heijden, G.W. & Bortvin, A. Optimized flow cytometry isolation of murine spermatocytes. Cytometry A 85, 556–565 (2014).

24. Bellani, M.A., Boateng, K.A., McLeod, D. & Camerini-Otero, R.D. The expression profile of the major mouse SPO11 isoforms indicates that SPO11beta introduces double strand breaks and suggests that SPO11alpha has an additional role in prophase in both spermatocytes and oocytes. Mol Cell Biol 30, 4391–4403 (2010).

25. Chakraborty, P. et al. LIN28A marks the spermatogonial progenitor population and regulates its cyclic expansion. Stem Cells 32, 860–873 (2014).

26. Kleene, K.C. & Flynn, J.F. Characterization of a cDNA clone encoding a basic protein, TP2, involved in chromatin condensation during spermiogenesis in the mouse. J Biol Chem 262, 17272–17277 (1987).

27. Zhou, Q. et al. Expression of stimulated by retinoic acid gene 8 (Stra8) in spermatogenic cells induced by retinoic acid: an in vivo study in vitamin A-sufficient postnatal murine testes. Biol Reprod 79, 35–42 (2008).

28. Tyanova, S. et al. The Perseus computational platform for comprehensive analysis of (prote)omics data. Nat. Methods 13, 731–740 (2016).

29. Poggio, T. & Girosi, F. Networks for approximation and learning. Proceedings of the IEEE 78, 1481–1497 (1990).

30. Cortes, C. & Vapnik, V. Support-Vector Networks. Machine Learning 20, 273–297 (1995).

31. Picard, R.R. & Cook, R.D. Cross-validation of regression models. Journal of the American Statistical Association 79, 575–583 (1984).

32. Kuleshov, M.V. et al. Enrichr: a comprehensive gene set enrichment analysis web server 2016 update. Nucleic Acids Res. 44, W90–W97 (2016).

33. Grootegoed, J.A., Jansen, R. & Van der Molen, H.J. The role of glucose, pyruvate and lactate in ATP production by rat spermatocytes and spermatids. Biochim Biophys Acta 767, 248–256 (1984).

34. Patel, M.S., Nemeria, N.S., Furey, W. & Jordan, F. The pyruvate dehydrogenase complexes: structure-based function and regulation. J Biol Chem 289, 16615–16623 (2014).

35. Takakubo, F. & Dahl, H.H. The expression pattern of the pyruvate dehydrogenase E1 alpha subunit genes during spermatogenesis in adult mouse. Exp Cell Res 199, 39–49 (1992).

36. Roeder, G.S. & Bailis, J.M. The pachytene checkpoint. Trends Genet 16, 395–403 (2000).

37. Ashley, T., Gaeth, A.P., Creemers, L.B., Hack, A.M. & de Rooij, D.G. Correlation of meiotic events in testis sections and microspreads of mouse spermatocytes relative to the mid-pachytene checkpoint. Chromosoma 113, 126–136 (2004).

38. Shao, B. et al. Unraveling the proteomic profile of mice testis during the initiation of meiosis. J Proteomics 120, 35–43 (2015).

39. Ahmed, E.A. & de Rooij, D.G. Staging of mouse seminiferous tubule cross-sections. Methods Mol Biol 558, 263–277 (2009).

40. Huang, T. et al. The histone modification reader ZCWPW1 links histone methylation to repair of PRDM9-induced meiotic double stand breaks. bioRxiv, 836023 (2019).

41. Li, M. et al. The histone modification reader ZCWPW1 is required for meiosis prophase I in male but not in female mice. Science advances 5, eaax1101 (2019).

42. Mahgoub, M. et al. Dual Histone Methyl Reader ZCWPW1 Facilitates Repair of Meiotic Double Strand Breaks. bioRxiv, 821603 (2019).

43. Wells, D. et al. ZCWPW1 is recruited to recombination hotspots by PRDM9, and is essential for meiotic double strand break repair. bioRxiv, 821678 (2019).

44. Peters, A.H., Plug, A.W., van Vugt, M.J. & de Boer, P. A drying-down technique for the spreading of mammalian meiocytes from the male and female germline. Chromosome Res 5, 66–68 (1997).

45. Cox, J. & Mann, M. MaxQuant enables high peptide identification rates, individualized p.p.b.-range mass accuracies and proteome-wide protein quantification. Nat. Biotechnol. 26, 1367–1372 (2008).

46. Charrad, M., Ghazzali, N., Boiteau, V. & Niknafs, A. NbClust: An R Package for Determining the Relevant Number of Clusters in a Data Set. 2014 61, 36 %J Journal of Statistical Software (2014).

47. Huang, D.W., Sherman, B.T. & Lempicki, R.A. Systematic and integrative analysis of large gene lists using DAVID bioinformatics resources. Nat. Protoc. 4, 44–57 (2009).

48. Waskom, M. et al. mwaskom/seaborn: v0. 9.0 (July 2018). Zenodo. (2018).

49. Breiman, L. Bagging predictors. Machine learning 24, 123–140 (1996).

50. Wang, H. et al. One-Step Generation of Mice Carrying Mutations in Multiple Genes by CRISPR/Cas-Mediated Genome Engineering. Cell 153, 910–918 (2013-05).

51. Yang, H. et al. One-step generation of mice carrying reporter and conditional alleles by CRISPR/Cas-mediated genome engineering. Cell 154, 1370–1379 (2013).

52. Yang, H., Wang, H. & Jaenisch, R. Generating genetically modified mice using CRISPR/Cas-mediated genome engineering. Nat Protoc 9, 1956–1968 (2014).

53. Perez-Riverol, Y. et al. The PRIDE database and related tools and resources in 2019: improving support for quantification data. Nucleic Acids Res 47, D442–D450 (2019).

