## Supplementary Note for "Prediction of meiosis-essential genes based on dynamic proteomes responsive to spermatogenesis"

**Supplementary Notes:**

**Regularized radial basis function network (Regularized RBF Network)**

Regularized radial basis function network[29] was applied as one type of sub-classifier for meiosis-essential protein prediction. The structure of this network is shown below.

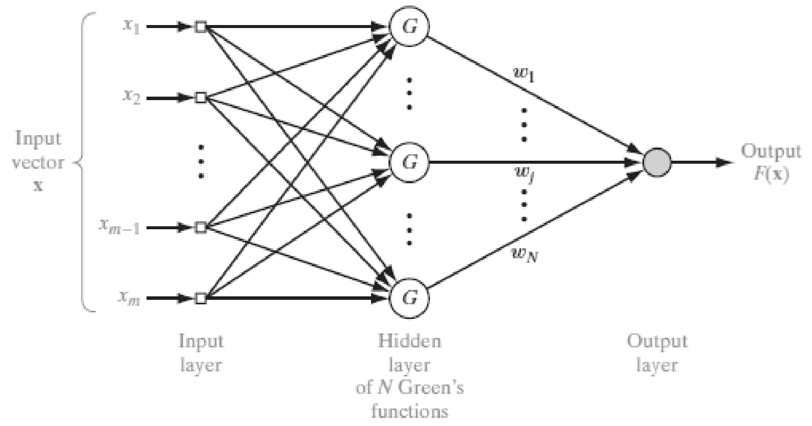


In this paper, $m$=9 and $\boldsymbol{x_1}, \boldsymbol{x_2}, \cdots, \boldsymbol{x_m}$ represented protein abundance of nine types of germ cells. $N,$ the number of Green's functions in the hidden layer, is equal to the number of proteins in training set. Several regularization methods could be applied to avoid the overfitting problem caused by the noise and uncertainty of data. In this study, we applied Tikhonov's regularization method. Based on the method, the loss function of the network contains two terms shown below.

$$\mathcal{E}(F) = \mathcal{E}_s(F) + \lambda \mathcal{E}_c(F)$$

$$\mathcal{E}_s(F) = \frac{1}{2}\sum_{i=1}^{N}\left(d_i - F(\boldsymbol{x_i})\right)^2 \ and\ \mathcal{E}_c(F) = \frac{1}{2}\|\boldsymbol{D}F\|^2$$

where $\mathcal{E}_s(F)$ is the standard error term and $\mathcal{E}_c(F)$ is the regularizing term, $\lambda$ is the regularization parameter and $\boldsymbol{D}$ is a linear differential operator.

In this study,

$$G(\boldsymbol{x}, \boldsymbol{x_i}) = e^{-\frac{1}{2\sigma^2}\|x - x_i\|^2} \text{ is the Green functions}$$

$$\boldsymbol{D} = \sum_n \alpha_n^{\frac{1}{2}} \left( \frac{\partial}{\partial x_1} + \frac{\partial}{\partial x_2} + \cdots + \frac{\partial}{\partial x_m} \right)^n \text{ where } \alpha_n = \frac{\sigma^{2n}}{n!\, 2^n}$$

In this study, $\sigma = 1$ and we estimated the optimal choice of $\lambda$ at about $3\times10^{-4}$. To avoid an effect of extreme values of RBF-predicted scores, we applied $F^* = tanh(F)$ to constrain the output scores into$[-1,1]$.

**Naive Bayesian Model (NBM)**

A Naive Bayesian Model was applied as the second type of sub-classifier for meiosis-essential protein prediction. $\boldsymbol{x} = (x_1, x_2, \cdots, x_9)$ represented expression abundance of each protein in nine types of cells. Two different classes of data, meiosis-essential and non-essential, were denoted $C_1$ and $C_2$ respectively. This sub-classifier is supposed to estimate $P(C_1\,|\boldsymbol{x})$ and $P(C_2\,|\boldsymbol{x})$ based on Bayes formula.

Bayes formula claims that $P(C_k\,|\boldsymbol{x}) = \frac{\prod_{i=1}^{9} P(x_i|C_k\,)P(C_k)}{P(\boldsymbol{x})}$ where $k$=1 or 2. We used $P(x_i|C_k\,) = \frac{s_{ki}}{s_k}$ and $P(C_k) = \frac{s_k}{s}$ to estimate the corresponding probabilities, where $s_k$ is the number of $C_k$ data in the training set, $s$ is the number of data in the training set and $s_{ki}$ is the number of $C_k$ data in the training set which has component $x_i$. As the data in this research is continuous, we divided each component into 12 classes. $P(\boldsymbol{x})$ as the common denominator was not considered, since the $P(C_k\,|\boldsymbol{x})$ can be calculated by normalization.

**Support Vector Machine (SVM)**

Support Vector Machine (SVM) was applied as the third type of sub-classifier for meiosis-essential protein prediction[30]. The structure of the sub-classifier is shown below.

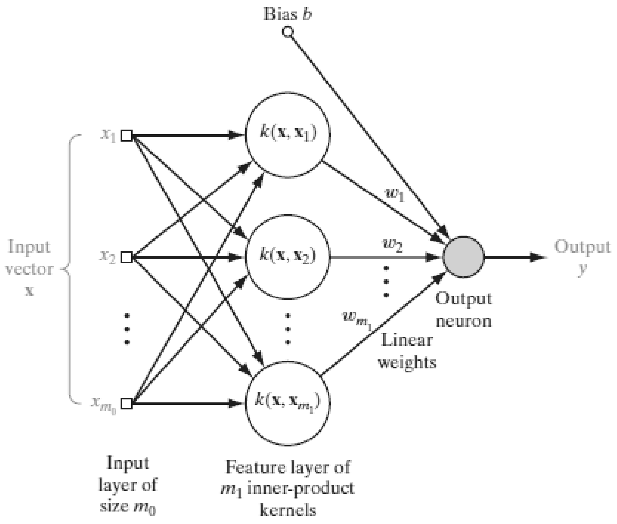


In this study, $m_0$=9 and $\boldsymbol{x_1}, \boldsymbol{x_2}, \cdots, \boldsymbol{x_{m_0}}$ represented expression abundance of each protein in nine types of cells, $\boldsymbol{x_1}, \boldsymbol{x_2}, \cdots, \boldsymbol{x_{m_1}}$ were expression abundance of all the proteins in the training set. We choose bias $b = 0$ and the kernel function $k(\boldsymbol{x}, \boldsymbol{x_i}) = e^{-\frac{1}{2\sigma^2}\|x - x_i\|^2}$ where $\sigma = 1$. In this study, we trained weights by solving the soft margin SVM problem shown below.

$$\min_{\boldsymbol{\omega}} \left( \frac{\lambda}{2} \|\boldsymbol{\omega}\|^2 + \frac{1}{m_1} \sum_{i=1}^{m_1} max\{0,\ 1 - d_i \langle \boldsymbol{\omega}, \boldsymbol{\psi}(\boldsymbol{x_i}) \rangle\} \right)$$

Where $d_i \in \{-1,1\}$ is the expected output, and $\langle \boldsymbol{\psi}(\boldsymbol{x}), \boldsymbol{\psi}(\boldsymbol{x_i}) \rangle = k(\boldsymbol{x}, \boldsymbol{x_i})$.

We applied the Stochastic Gradient Descent (SGD) method[53] to solve the problem and let $\lambda = 1$.

To avoid an effect of extreme values of SVM-predicted scores, we applied $y^* = tanh(y)$ to constrain the output scores into$[-1,1]$.
