## Supplementary Figures for "Prediction of meiosis-essential genes based on dynamic proteomes responsive to spermatogenesis"

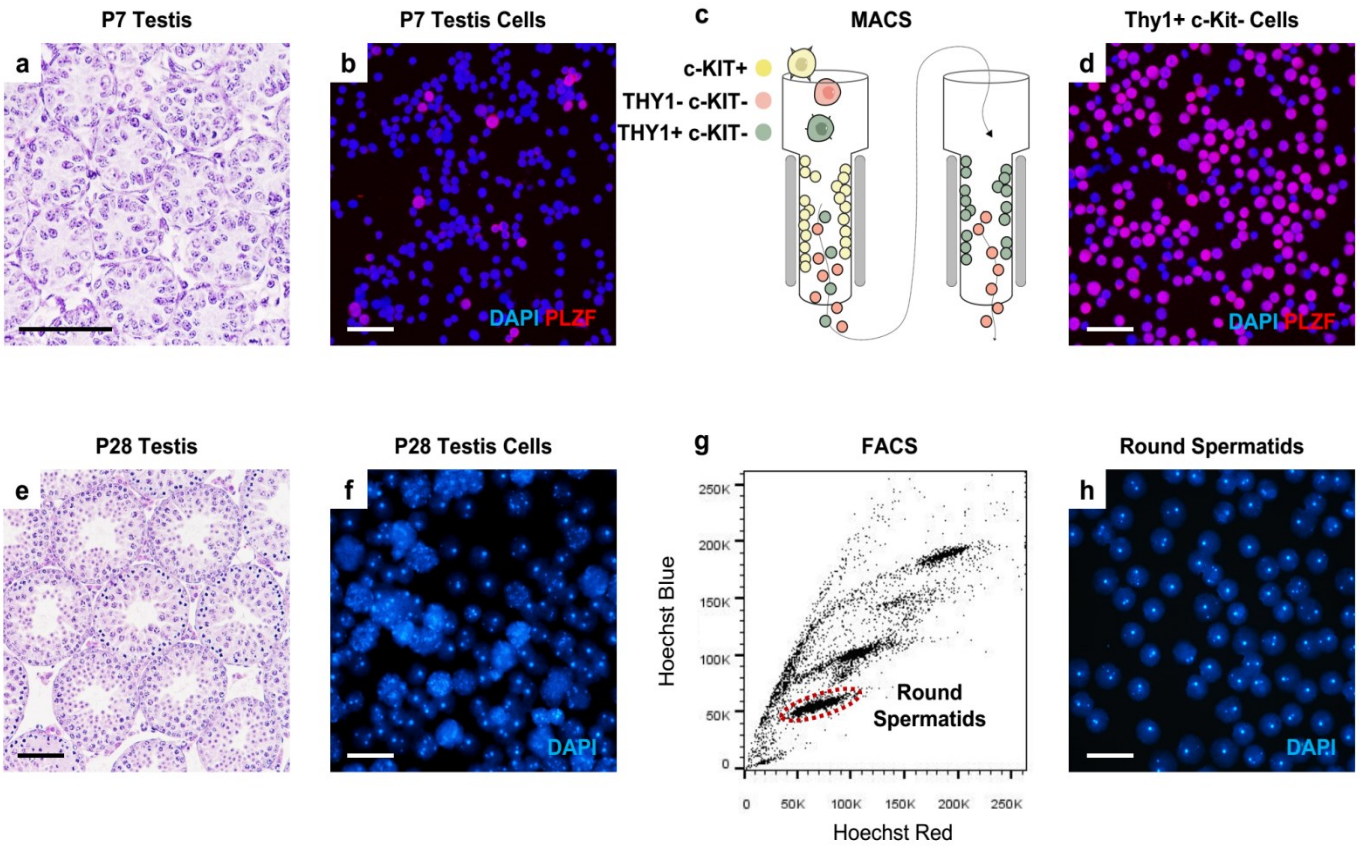


**Fig. S1 Isolation of undifferentiated spermatogonia (Aundiff) and round spermatids (RS)**. **a** Cross section of H&E stained testes from P7 mouse. **b** DAPI (blue) and anti-PLZF (red) immuno-staining of digested P7 testes cells before MACS purification. **c** Illustration of magnetic activated cell sorting (MACS) strategy to isolate THY1+ c-KIT- Aundiff. Differentiated spermatogonia and other testes somatic cells expressed c-KIT (c-KIT+ cells, shown as yellow cells), which were depleted first by binding to the anti-c-KIT antibody and magnetic columns. The unbound c-KIT- cells were subsequently separated by MACS to enrich the THY1+ c-KIT- cells (showed as green cells). **d** DAPI (blue) and anti-PLZF (red) immuno-staining of THY1+ c-KIT- cells after MACS purification. **e** Cross section of H&E stained testes of P28 mouse. **f** DAPI staining of digested P28 testes cells before FACS purification. **g** FACS plot of digested testes cells stained with Hoechst 33,342. Gate for sorting haploid RS is circled. **h** DAPI staining of RS after FACS purification. All scale bars = 50 µm.

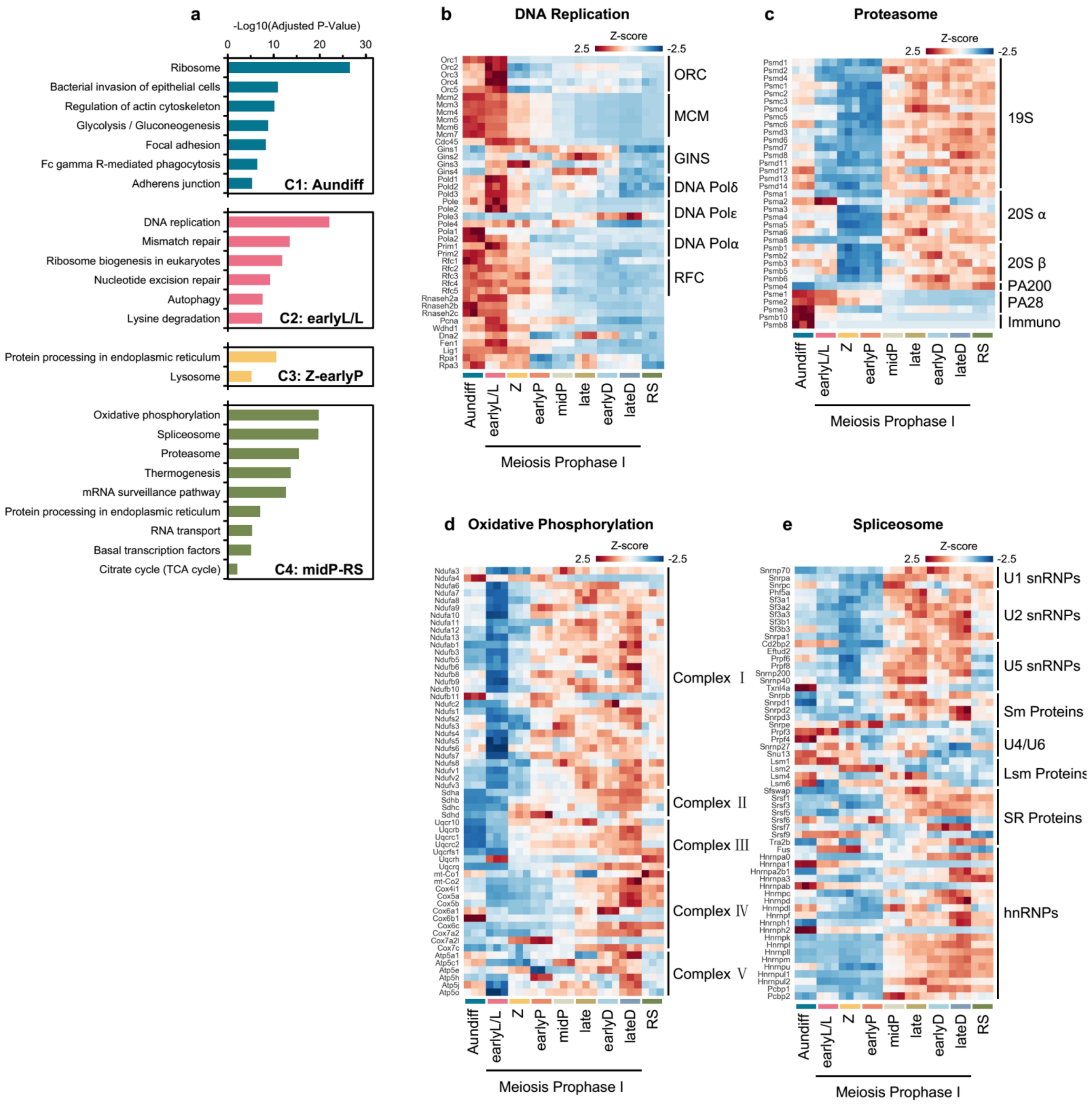


**Fig. S2 Enriched KEGG pathways for four DEP clusters. a** KEGG pathway analysis of the enriched functions for four DEP clusters. **b-e** Heatmap of dynamic abundance of the proteins involved in DNA replication (**b**), proteasome (**c**), oxidative phosphorylation (**d**) and spliceosome (**e**). Color key represents the Z-score of relative protein abundance.

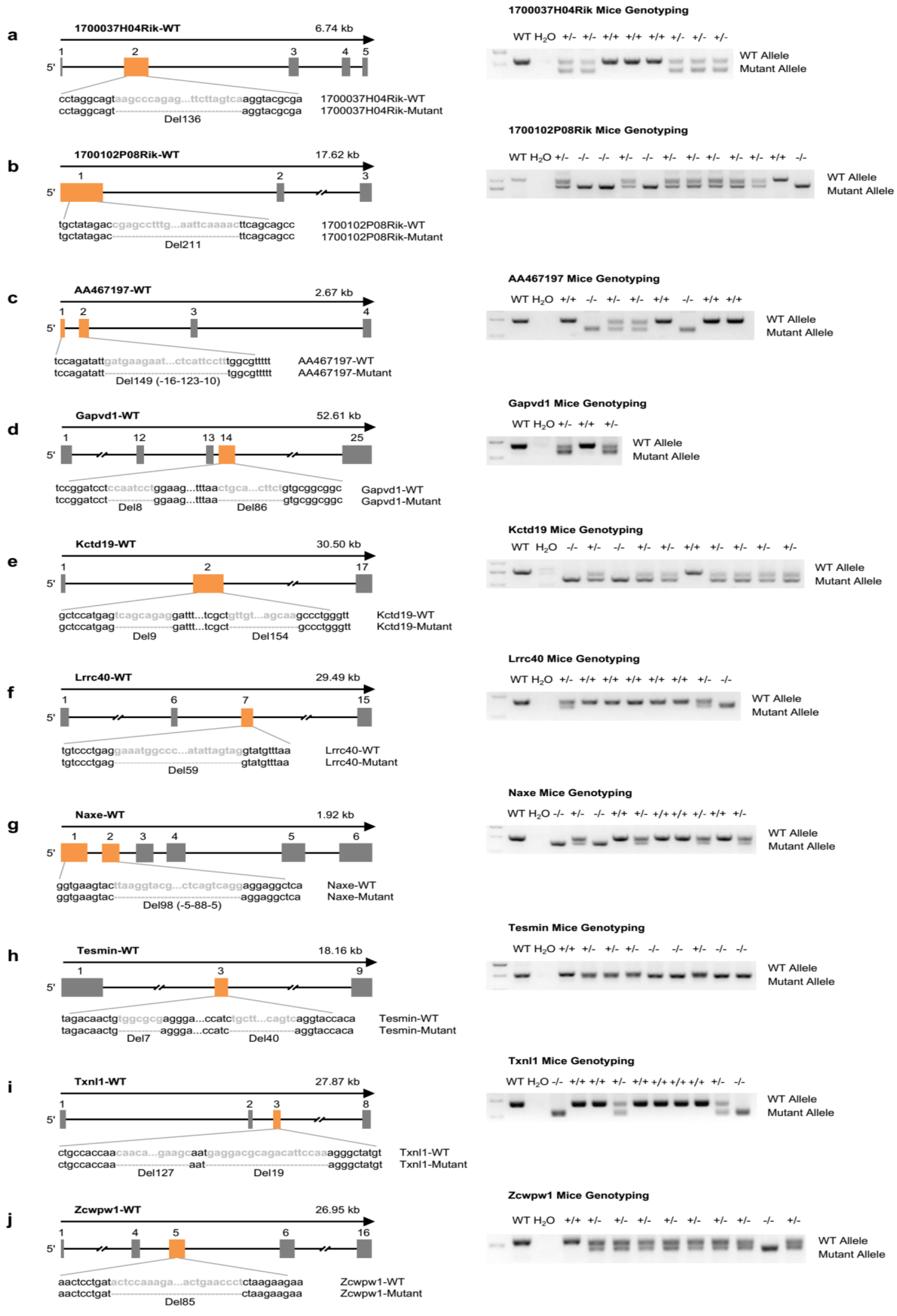


**Fig. S3 Construction and genotyping of ten knock out mice of the top ten RBF-ranked candidates. a-j** Strategy to construct knockout mice of the top 10 RBF-ranked candidates (left panel) and the genotyping of knockout mice verified by PCR (right panel).

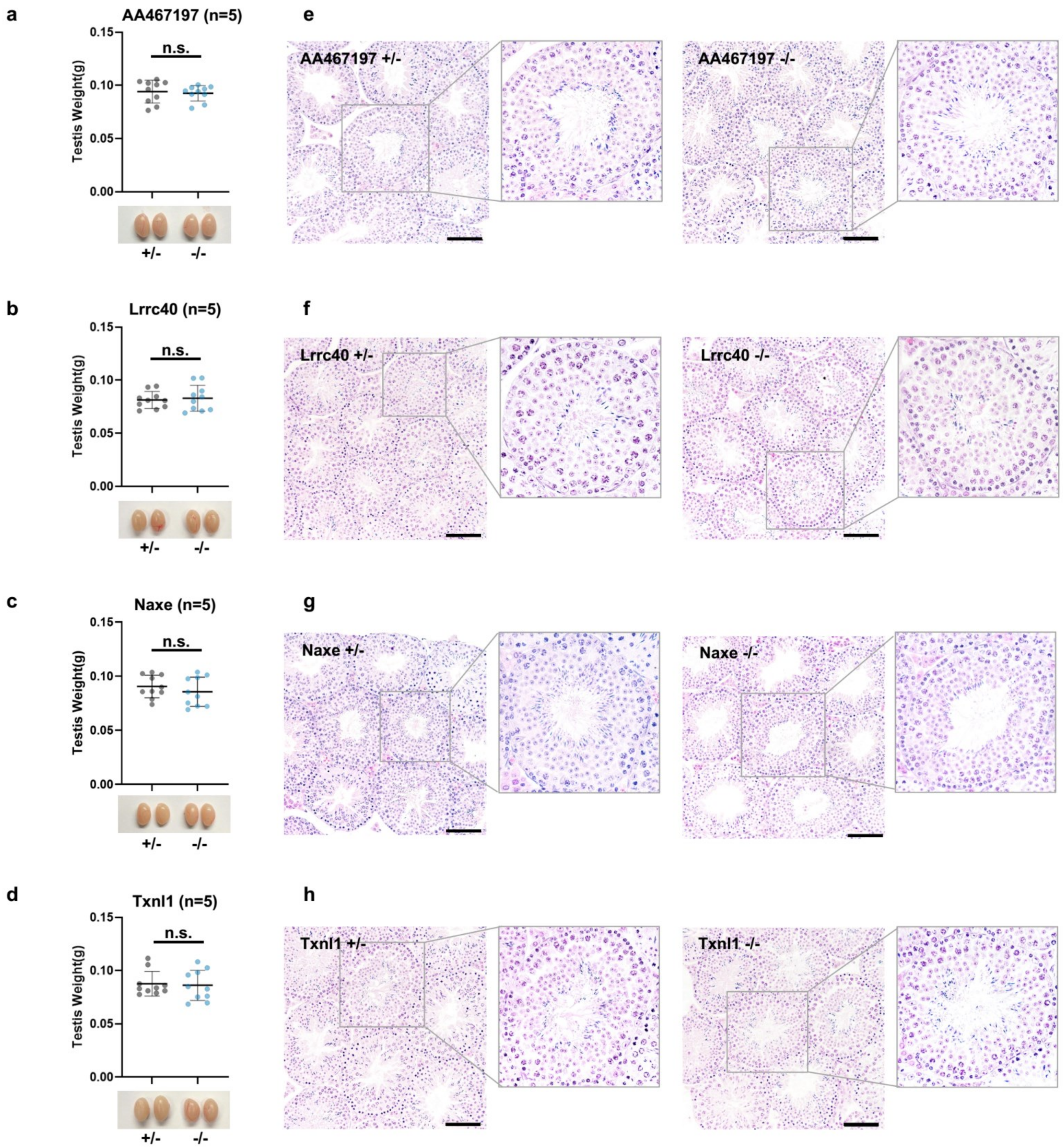


**Fig. S4 Phenotypic validation of the meiosis non-essential proteins predicted by RBF**. **a-d** Comparison of testis weights derived from 8-week-old Txnl1$^{+/-}$ and Txnl1$^{-/-}$ mice (**a**), AA467197$^{+/-}$ and AA467197$^{-/-}$ mice (**b**), Lrrc40$^{+/-}$ and Lrrc40$^{-/-}$ mice (**c**), and Naxe$^{+/-}$ and Naxe$^{-/-}$ mice (**d**). n.s. means no significance in unpaired two-tailed *t*-test. **e-h** Cross-sections of H&E stained seminiferous tubules from the heterozygous and homozygous knockout mice of the four genes above, insets denote the specific seminiferous tubule under higher magnification. All scale bars = 50 µm.
